# Recording the age of RNA with deamination

**DOI:** 10.1101/2020.02.08.939983

**Authors:** Samuel G. Rodriques, Linlin M. Chen, Sophia Liu, Ellen D. Zhong, Joseph R. Scherrer, Edward S. Boyden, Fei Chen

**Author notes:** These authors contributed equally to this work (Samuel G. Rodriques, Linlin M. Chen, Edward S. Boyden, Fei Chen).

## Abstract

Transcriptional programs implemented by cells often consist of complex temporal features, but current approaches to single-cell RNA sequencing only provide limited information about the dynamics of gene expression. Here, we present RNA timestamps, a method for inferring the age of individual RNAs in a sequencing-based readout by leveraging RNA editing. Timestamped RNAs include a RNA reporter motif that accumulates A to I edits over time, allowing the age of the RNA to be inferred with hour-scale accuracy. By combining observations of multiple timestamped RNAs driven by the same promoter, we are able to infer when in the past the promoter was active. We demonstrate that the system can infer the presence and timing of multiple past transcriptional events, with no prior knowledge. Finally, we apply this method to cluster single cells according to the times at which a particular transcriptional program was activated. RNA timestamps thus suggest a new approach for incorporating temporal information into RNA sequencing workflows.

## Main

Rapid progress in RNA-sequencing technologies, and, in particular, single-cell RNA-seq (scRNA-seq), has enabled valuable insights and molecular characterization of complex tissues. However, since sequencing is a destructive measurement, these approaches only provide an instantaneous snapshot of cellular state, and comparatively little work has been done to examine how cell states arise from complex transcriptional dynamics over timescales of hours. Current approaches to add a temporal dimension to RNA sequencing through metabolic labeling of newly synthesized RNAs (*1*–*4*), or computational inference from the abundance of unspliced transcripts (*5*), only provide the derivative of expression level or information about expression at a single timepoint, so transcriptional histories must still be reconstructed by observations of many cells. To overcome these challenges, we asked whether it might be possible to estimate the age of *each individual RNA*, eventually allowing us to build up a picture of the transcriptional history of a single cell from the ensemble of RNAs present in the cell.

To that end, we designed a recorder RNA motif, referred to as an RNA timestamp, that can report its own age via the gradual accumulation of A to I edits caused by an engineered version of the human Adenosine Deaminase Acting on RNA 2 catalytic domain (ADAR2cd, **Fig. 1A**). This catalytic domain has previously been shown to targetable to RNAs via fusion to exogenous RNA binding domains, with minimal off-target activity to untargeted RNAs (*6*–*10*). The timestamps are adenosine-rich editing arrays that are designed to be favored substrates of the ADAR enzyme (*11*–*13*) (**Fig. 1B**). Edits in this region can subsequently be identified as A to G mutations in high throughput sequencing of the timestamps. ADAR2cd is specifically targeted to MS2 binding sites in the editing region of the timestamps through a fusion with the MS2 Capsid Protein (MCP)(*14*). We screened multiple RNA and ADAR variants, and selected a pair with an editing timescale on the order on the order of hours, a timescale relevant for endogenous transcriptional activity (**Supp. Fig. 1**).

**Figure 1:**
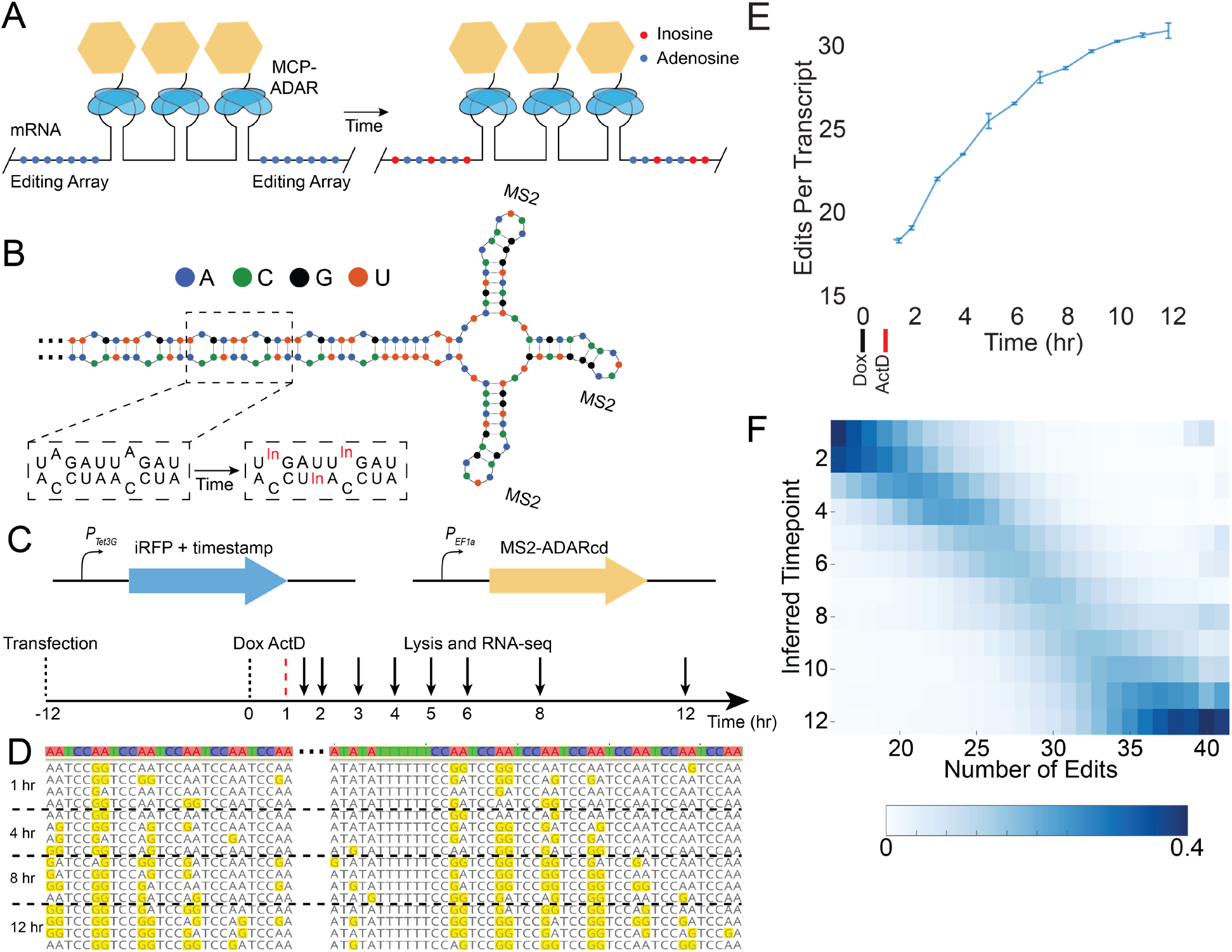
Encoding of temporal information through RNA edits. (**A**) Timestamps consist of editing arrays of adenosines (blue dots) and several MS2 step loops in the 3’ UTR of an mRNA. In the presence of an MCP-ADAR fusion (MCP, blue ellipses, ADAR, yellow hexagon), timestamps are edited over time by catalytic conversion of adenosine to inosine (red dots). (**B**) The structure of a portion of one timestamp, showing MS2 stem loops and the repetitive, double-stranded RNA motif that serves as the editing substrate. (**C**) Schematic representation of the Tet-responsive timestamped mRNA system and experimental timeline. (**D**) Examples of timestamps edited for different amounts of time, showing the accumulation of A>I edits. (**E**) The mean number of edits per timestamp observed after editing for different amounts of time. Error bars show standard deviation over 3 biological replicates. (**F**) The number of edits per timestamp is shown for different specified ages. Columns are normalized to 1; each column can be interpreted as a probability distribution of when a timestamped RNA with a given number of edits is likely to have been made.

In order to calibrate our system, we incubated HEK293T cells expressing the ADAR variant and a timestamped RNA under the control of the tetracycline response element (TRE) in medium containing doxycycline for one hour. The timestamp was placed in the 3’ UTR of a mRNA encoding a fluorescent protein (*15*) that was optimized to prevent degradation over time (**Supp. Fig. 2**), for purposes of calibration. We subsequently added actinomycin D, an RNA transcription inhibitor (*16*), and then lysed the cells after a variable amount of time before sequencing the timestamps (**Fig. 1C**). Each of these experiments provided us with a population of RNAs with a known age (± 30 minutes), and we observed that the timestamps became progressively more heavily edited over time (**Fig. 1D**). The mean number of edits per timestamp at any given timepoint was highly reproducible between biological replicates (**Fig. 1E**), and the accumulation of edits over time could be modeled accurately using a Poisson-Binomial model (**Supp. Fig. 3**). Using the empirical distribution of the number of edits per timestamp, we found that it was possible to estimate the age of a timestamped mRNA transcript purely from the total number of edits on the timestamp with a 95% confidence interval of 2.7 ± 0.4 hours (mean ± stdev, N=26 bins of timestamp edits, **Fig. 1F**).

Since cells will often contain many copies of a given RNA transcript, we reasoned that we might be able to infer the underlying transcriptional dynamics of a specific promoter by analyzing the ensemble of timestamped RNAs produced by the promoter. To test this, we built an algorithm that infers the transcriptional program that is most consistent with a particular set of timestamps (see Methods). The algorithm accomplishes this by using gradient descent to identify a convex combination of single-hour editing distributions that minimizes the L2 norm between the observed editing distribution and the editing distribution associated with the convex combination (**Fig. 2A**). The algorithm makes no assumptions about the underlying transcriptional program that generated the observed timestamps. We applied the algorithm to sets of timestamped RNAs that were all chosen randomly from the same timepoint, and found that it could infer both that the sets came from a single timepoint, and which timepoint they came from, with accuracy that increased monotonically with the number of RNAs in the set (**Fig. 2B**). Consistent with our hypothesis that temporal estimates based on multiple RNAs would be more accurate than temporal estimates based on a single RNA, the temporal error associated with a set of only 4 RNAs was 1.49 ± 0.42 h (mean ± std, N=12 timepoints, see Methods), and declined monotonically with increasing numbers of RNAs (**Fig. 2C**). For sets of at least 50 RNAs, chosen because it is consistent with the number of RNAs one might find in an individual cell in response to a stimulus (*17*), the algorithm reproduced weight distributions that clearly resembled the single-hour transcriptional programs from which the RNAs were derived (**Fig. 2D**).

**Figure 2:**
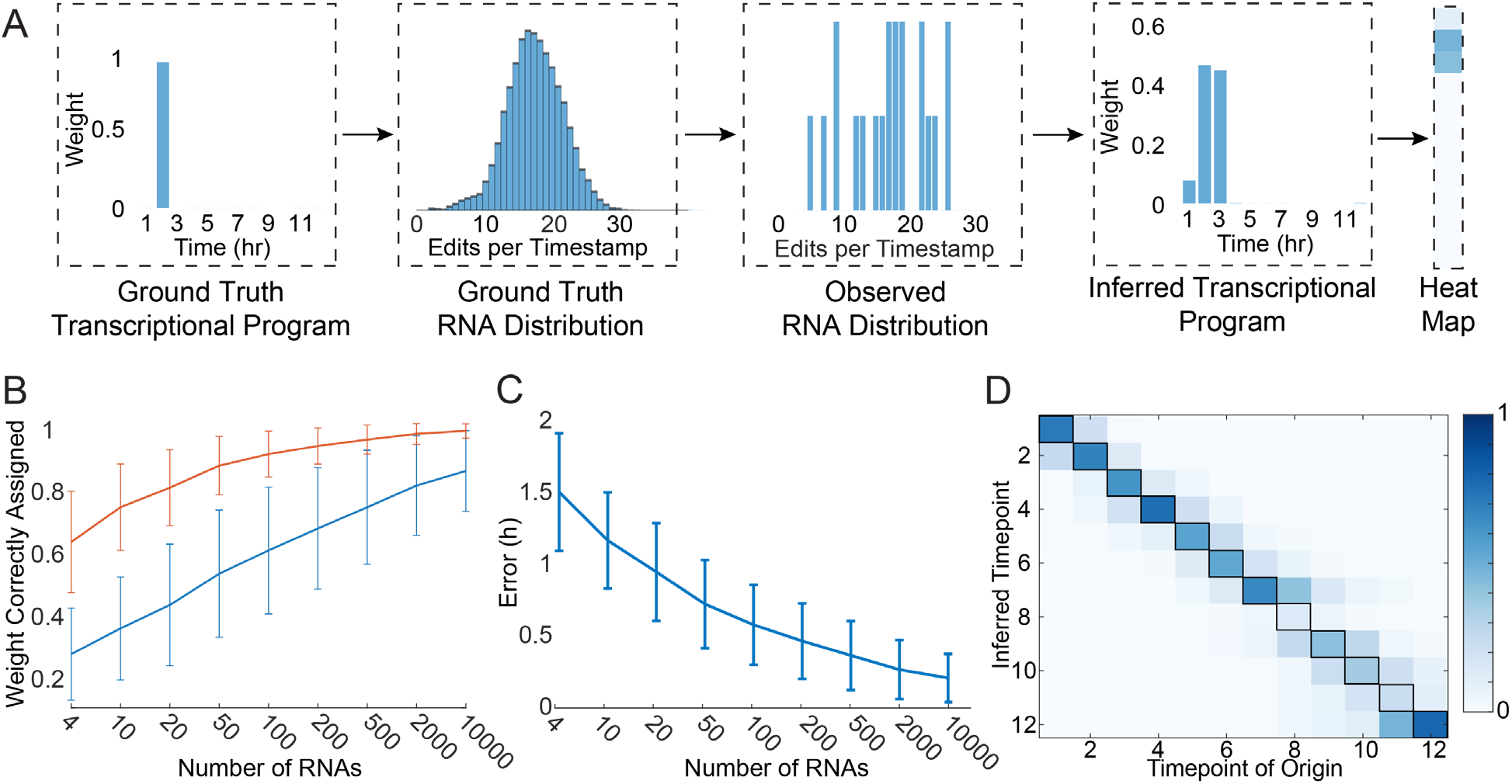
Timestamped RNAs can reveal temporal transcription programs. (**A**) Schematic of the gradient descent algorithm. RNAs are generated by a ground truth underlying transcriptional program (e.g. a transcriptional event that occurred two hours prior to lysis, left), which corresponds to a ground truth distribution of the number of edits per timestamp (middle left). In any individual experiment, one observes a set of timestamped RNAs sampled from the true underlying distribution (middle right). The gradient descent algorithm produces an inferred transcriptional program (right) that best approximates the observed RNA distribution under the L2 norm. Transcriptional programs can then be represented as columns in a heat map (far right). (**B**) For sets of timestamped RNAs drawn from a single timepoint, the fraction assigned by the algorithm to the correct timepoint (blue) or the correct three-hour window (orange) as a function of the number of RNAs in the set. Error bars show std (N=12 timepoints). (**C**) The temporal error associated with the reconstructed transcriptional programs, as a function of the number of RNAs in the set. See Methods. (**D**) The average weight assigned by the gradient descent algorithm to each timepoint (Y axis) is shown as a function of the ground truth timepoint of origin (X axis), for sets of 50 RNAs. Columns sum to 1.

Since each RNA is individually timestamped, we reasoned that the algorithm might be able to identify if a given set of timestamped RNA was produced by multiple, temporally distinct transcriptional pulses. To test this, we computationally generated mixtures of RNAs drawn half from the 1-hour timepoint and half from the 5-hour timepoint, and applied the gradient descent algorithm to infer the transcriptional program most likely to have generated the mixtures (**Fig. 3A**). Even with only 10 RNAs, we were able to identify the presence of two peaks in the transcriptional program 51.3% ± 7.6% of the time, and this increased to 86% ± 13% with 200 RNAs (mean ± std, N=3 replicates, **Fig. 3B**, see Methods). To validate these predictions experimentally, we transfected cells with barcoded RNAs driven by a doxycycline-sensitive promoter and a light-sensitive transcription factor (*18*). Timestamped RNAs associated with each promoter could be identified by promoter-specific barcodes, allowing for independent reconstruction of light- and doxycycline-induced timestamp editing distributions (**Supp. Fig. 4**). Independently stimulating the cells with each promoter at two different times (at 1 and 5 hours) led to an editing distribution with two clear peaks corresponding to the two transcriptional events (**Fig. 3C**). We then subsampled the timestamped RNAs from this experiment (**Fig. 3D**) and ran the gradient descent algorithm to generate putative transcriptional programs (**Fig. 3E**). Even though this was a more challenging task, since the RNAs were not chosen evenly from the two peaks, the results we obtained were very similar to the results obtained in our simulations (**Fig. 3F**). With 10 RNAs, two peaks could be distinguished 44.5% ± 0.7% of the time, and this increased to 90.5% ± 0.7% with 200 RNAs (mean ± std, N=2 replicates, **Fig. 3G**).

**Figure 3:**
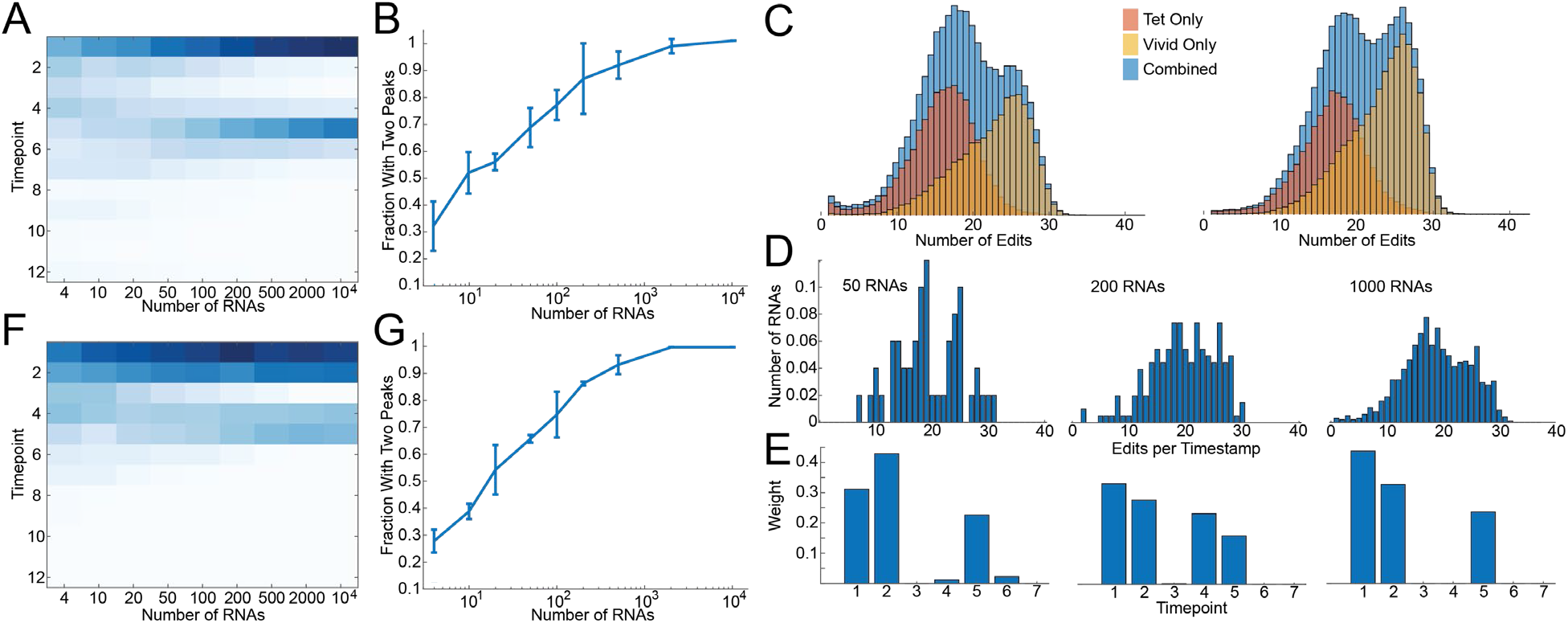
Identification of temporally separated transcriptional events. (**A**) The average weight assigned by the gradient descent algorithm to each timepoint for sets of timestamped RNAs that were drawn randomly (in equal proportion) from the 1 and 5-hour timepoints as a function of number of RNA drawn. Columns sum to 1. (**B**) The percentage of sets of RNA in which two peaks can be identified by the gradient descent algorithm (see Methods). Error bars show std, N=3 datasets from biological replicates. (**C**) Histograms of the number of edits per timestamp are shown for cells that were induced separately with a light-sensitive promoter and a doxycycline-sensitive promoter. Red shows the distribution of edits per RNA for RNAs transcribed off the doxycycline promoter, while yellow shows the light-sensitive promoter. Blue shows the combined distribution. Columns show two biological replicates. (**D**) Distributions of edits per timestamp for sets of RNAs chosen randomly from the blue distribution in (C). (E) The weight distribution inferred by the gradient descent algorithm for the set of RNAs shown in (D). Same as (A), but for sets of RNA drawn from the experimental distributions in (C). (**G**) Same as (B), but for sets of RNA drawn from the experimental distributions in (C). Error bars show std, N=2 biological replicates.

Given the performance of the timestamp system with relatively few RNAs, we asked whether it could be applied to determine the timing of transcriptional events in individual cells. We transfected cells with timestamped RNAs under the control of TRE, induced with doxycycline, and then induced the cells according to three different protocols, including a many-hour induction protocol and two single-hour induction protocols (see Methods). Individual cells were then sorted into wells of a 96 well plate, and we subsequently performed single-cell timestamp sequencing (**Fig. 4A**). Analysis of the editing histograms yielded a mean induction time associated with each cell (see Methods). The estimation error in the temporal estimate for the single cells in the single-hour induction protocols were 0.2hr +/− 0.8hr (mean +/− std, N=27) for one condition and 0.5hr +/− 1.0hr (mean +/− std, N=19) for the other (**Fig. 4B**, see Methods). Based on these temporal estimates, we were able to order the cells according to the timing of transcriptional events, obtaining only 5 transpositions out of 72 cells, an accuracy rate of 86% (**Fig. 4C**).

**Figure 4:**
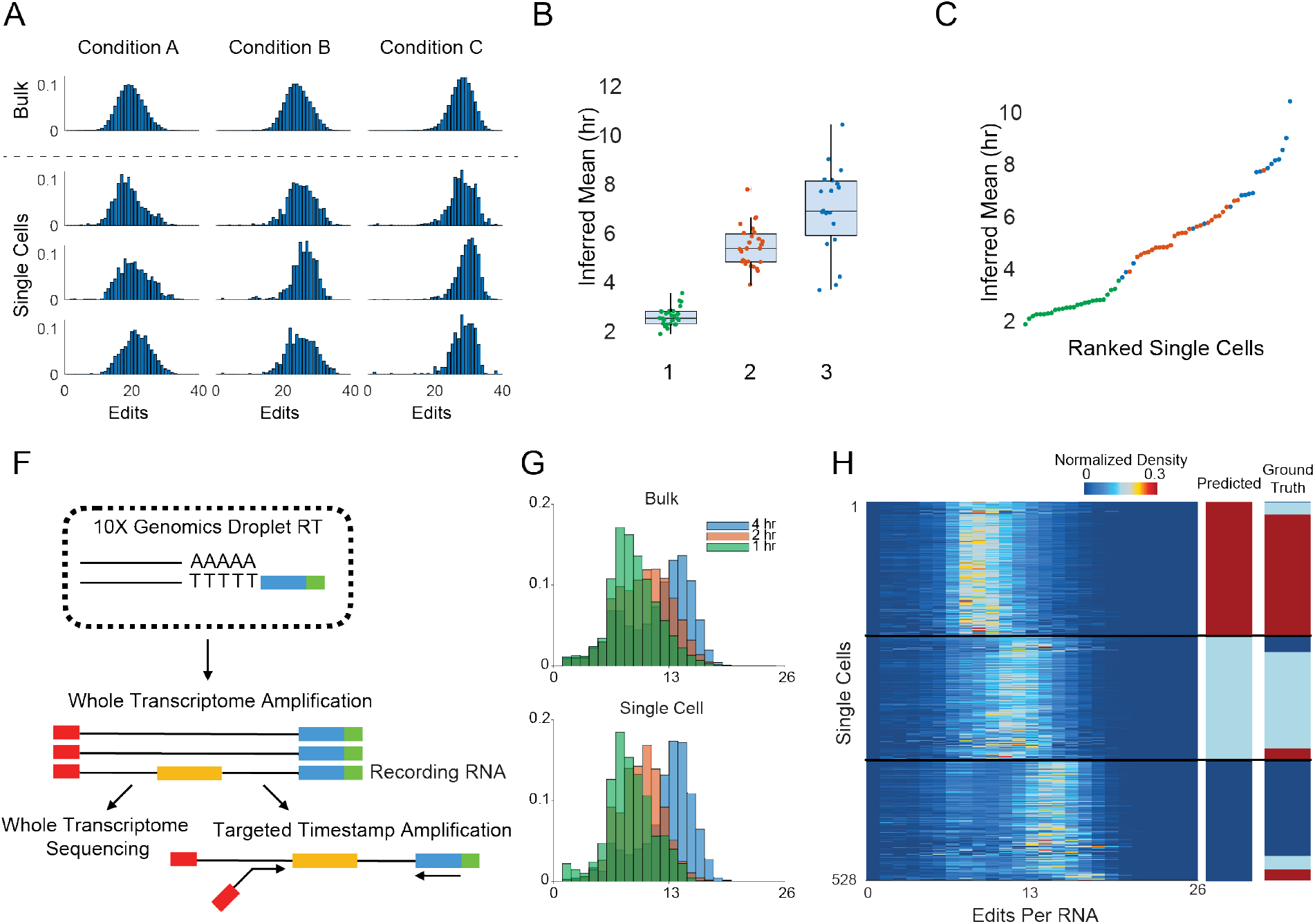
Timestamps can reveal transcriptional programs in single cells. All editing histograms are normalized to sum to 1. (**A**) Editing histograms for bulk conditions 1 through 3 (top) and randomly chosen single cells (bottom). (**B**) Predicted induction times for all single cells in the experiment, calculated as the center of mass of the inferred weight distributions (N=24 for condition 1, N=27 for condition 2, N=19 for condition 3). (**C**) The predicted induction time for each single cell, ranked from least to greatest. Green dots correspond to condition 1; red dots correspond to condition 2; blue dots correspond to condition 3. (**D**) Schematic of capture and sequencing of timestamps with 10x Single Cell sequencing in conjunction with transcriptomic profiling. (**E**) Top: Editing histograms for bulk sequencing of 1-hour induction events 1, 2 and 4 hours prior to lysis. Bottom: Three representative single cells for 1, 2, and 4 hour induction conditions (rows 50, 250 and 450 from H). (**F**) Heatmap of distribution of edits per cell for 528 cells. Cells are clustered by k-means according to their editing distribution into 3 clusters corresponding to predicted induction. “Ground truth” column shows grouping of cells by condition, determined using a condition barcode on the timestamp reads (Red: 1 hr, Light Blue: 2 hr, Dark Blue, 4 hr).

Recent advances in droplet-based barcoding technologies have enabled the scalable molecular profiling of cells and tissues at the single cell level (*19*, *20*). Thus, we asked if we could read out timestamps with 10x Single Cell 3’ RNA sequencing in combination with paired transcriptomic information. To do so, we developed a molecular biology workflow which enables targeted amplification of timestamps from single-cell barcoded whole transcriptome amplification products (see Methods; **Fig. 4D**). This allows for both the timestamps and the cell barcode to be read out via paired-end Illumina sequencing. We transfected HEK cells with the barcoded TRE timestamp systems and induced with doxycycline 1, 2 or 4 hours before 10x droplet-based sequencing. Single cell recRNA recordings measured via 10x sequencing is qualitatively similar to bulk sequencing from the same conditions (**Fig. 4E**). After filtering for high quality cells, we performed k-means clustering on the timestamps, and found three distinct clusters (528 cells, ~68% of cells) corresponding to the 1, 2 and 4 hour inductions (the other 32% corresponded to cells with few edits, potentially without ADAR). Examining these 528 cells, we found that 437/538 (~81%) of these cells were correctly assigned to an induction condition when compared to ground truth (**Fig. 4F**). These results demonstrate that timestamps can be read out in concert with high throughput single cell droplet-based methods.

Here, we have reported that the continuous enzymatic activity of adenosine deaminases acting on RNA can allow for direct inference of the age of individual RNAs, and that statistical methods taking advantage of this phenomenon can reveal the temporal aspect of transcriptional dynamics in individual cells. This observation may find widespread utility: for example, the demonstrated ability to order cells by transcriptional response could have great utility for studying the diversity of responses to cellular perturbations (*21*, *22*). Using the same concept, alternative systems could be designed that record other kinds of signals besides transcription. For example, by using alternative dimerization systems(*23*, *24*) to link ADAR to constitutively expressed timestamps in a stimulus-specific manner, it may be possible to construct timestamp systems that report on the timing of other kinds of cellular events, such as calcium or other signaling molecules. Moreover, we suspect that timestamps could readily be added to endogenous RNAs, or that a similar concept could be used, for example by observing edits in the poly(A) tail, or by examining adenosine-rich parts of the transcript. Together, these observations suggest that RNA timestamps provide a scalable and extensible approach for recording the temporal activity of cells.

## Acknowledgments

We acknowledge Noah Jakimo, Asmamaw T. Wassie, Jonathan Gootenberg, and Omar Abuddayeh for helpful discussions. Plasmids containing ADAR2 mutants were generously provided by Jonathan Gootenberg and Omar Abuddayeh. Plasmids containing the iRFP gene were generously provided by Changyang Linghu. Neuron culture was supplied by Demian Park. We acknowledge Yingxi Lin and Xiaochen Sun for help with neuron induction experiments. All data will be available online, and all plasmids are deposited on Addgene.

F.C. acknowledges funding from 1DP5OD024583, NIH Directors Early independence award and Schmidt Fellows Program at Broad Institute. E.S.B. acknowledges funding by John Doerr, the Open Philanthropy Project, NIH 1R01MH114031, the HHMI-Simons Faculty Scholars Program, U. S. Army Research Laboratory and the U. S. Army Research Office under contract/grant number W911NF1510548, NIH 1RM1HG008525, NIH 2R01DA029639, NIH 1R01MH103910, NIH Director’s Pioneer Award 1DP1NS087724, and the MIT Media Lab. E.S.B. also acknowledges Lisa Yang as a supporter of his lab. S.G.R. acknowledges funding through the Hertz Graduate Fellowship and the National Science Foundation Graduate Research Fellowship Program (award #1122374). J.S. acknowledges funding through the Hertz Graduate Fellowship. S.L. acknowledges funding through the Molecular Biophysics Training Grant, NIH/NIGMS T32 GM008313. E.D.Z. acknowledges funding through the National Science Foundation Graduate Research Fellowship Program (award #1122374) and through the Computational and Systems Biology training grant, T32 GM087237.

## Author Contributions

S.G.R., F.C., and E.S.B. conceived of strategies for the design of the RNA timestamps. S.G.R., L.M.C., and J.S. conceived of and implemented the design of the reporter RNAs. S.G.R., L.M.C., F.C., and S.L. validated and characterized the timestamp system in cells. E.D.Z. conceived of and implemented the Poisson binomial model in Figure 2. S.G.R. and E.D.Z. conceived of and implemented the gradient descent model in Figure 3. S.G.R. and F.C. analyzed the data. S.G.R., F.C. and E.S.B. wrote the manuscript.

## Methods

### Cloning

All plasmids were constructed either using restriction cloning using restriction enzymes from New England Biosciences and the NEB Quick Ligation kit (M2200L), or using the In-Fusion HD cloning enzyme mix (Clontech, 638911). Plasmids were grown in E.Cloni 10G Chemically Competent Cells (Lucigen, 60107-1) and were verified by Sanger sequencing (Eton biosciences). All plasmids are deposited on Addgene.

Due to high repetition present in the RNA editing templates, inserts for plasmids 76, 147, 148, 149, and 187 (see Table S2) were ordered as sense and antisense ultramer oligonucleotides, which were annealed to each other prior to cloning. Plasmid 76 was cloned by inserting RNA templates (A_Short, B_Short, C, D, E) into the 3’ UTR of an iRFP transcript expressed under a UbC promoter in a second generation lentivirus backbone using SphI and ClaI. Subsequently, this plasmid was modified by the addition of a flavivirus xrRNA in the 5’ UTR. Templates A_Short and B_Short were then extended by inserting another pair of annealed ultramers on the 5’ side of A_Short and B_Short using SphI and MluI. The resulting templates are designated A and B. To generate plasmids 147, 148, 149, and 183 (as used in the paper), templates A and B were then moved into different backbones and different promoters by restriction cloning, or by Gibson assembly with PCR amplification of the timestamp template region. Template A is used throughout the paper, and Template B is shown in Supp. Fig. 1 for comparison.

### RNA Purification, Library Preparation, and Sequencing

All cell cultures were lysed with 600uL of buffer RLT Plus from the Qiagen RNEasy Plus Mini Kit (Qiagen, 74136), and were pipetted up and down vigorously to homogenize. RNA was then purified using the Qiagen RNEasy Plus Mini kit, following the instructions from the manufacturer. Subsequently, 11uL of purified RNA was reverse transcribed using Superscript IV (Thermofisher, 18090050) and a barcoded version of SGR-174 (see Table S2), following the protocol from the manufacturer. Reverse transcription reactions were then purified using Agencourt Ampure XP beads at a 1:1 dilution (Beckman-Coulter, A63881). Some portion of the eluent, typically 25%, was then PCRed using P5 and a barcoded version of SGR-176 (see Table S2) the Q5 Hot Start High Fidelity 2x Master Mix (NEB, M0492L) with the following settings: 30s of 98C denaturation; then 25-30 cycles of 10s denaturation at 98C, 20s annealing at 70C, and then 25s extension at 72C. Neuron lysates were typically PCRed for 30 cycles, while HEK cell lysates were typically PCRed for 25 cycles. PCR reactions were then pooled and run on a gel, and a 400bp band was extracted using the NucleoSpin PCR Cleanup Kit (Macherey-Nagel, 740609.250). The concentration of DNA in the resulting eluent was determined via a Qubit 2 fluorometer (Thermofisher), and was then adjusted to 4nM for sequencing. The read structure is shown in Supp. Fig. 6.

Sequencing was performed using NextSeq Mid Output 300 cycle kit (Illumina, FC-404-2004), Miseq 300 cycle v2 kits (MS-102-2002), or Miseq 600 cycle v3 kits (MS-102-3003), with at least 80bp read 1 and 185bp read 2, with 8bp index 1 and 15bp index 2.

### HEK and 3T3 cell culture

Except in the case of the single cell experiments, HEK293FT and 3T3 cells were plated in 24 well plates. Cells were grown in DMEM (Thermofisher, 10566016), supplemented with Penicillin/Streptomycin (Thermofisher, 15140122) and 10% certified Tet-system approved FBS (Clontech, 631101). Transfections were performed using the TransIT-X2 system (Mirus, MIR 6000), following the manufacturer’s instructions.

For doxycycline experiments, HEK and 3T3 cells in 24 well plates were transfected with 300ng of plasmid 147 or 148, 100ng of pCMV Tet3G from the Tet-on 3G system (Clontech, 631168), and 100ng of plasmids 116v1, 116v5, or 116v6. In the experiments for Figures 1, and Supp. Figs. 1 and 3, they were transfected with both 147 and 148, and received 150ng of each plasmid. At least 12 hours after transfection, cells were stimulated by adding doxycycline to a final concentration of 1ug/mL, followed by gentle mixing or swirling of the plate. Subsequently, transcription was halted by adding Actinomycin D to a final concentration of 1ug/mL in the same medium. After waiting for the experimental time period, cells were lysed using Buffer RLT Plus and libraries were prepared as described above.

For experiments using the Vivid promoter, including Fig. 3C-G and Supp. Fig. 4, 3T3s were transfected with 300ng of plasmid 149, 100ng of pCMV Tet3G, and 100ng of plasmid 116v5. For conditions in which cells were transfected with both plasmid 147 and plasmid 149, they received 150ng of each plasmid. Cells were stimulated with a blue LED (Thor Labs, M455L2) with a total power of 200uW/cm^2. The LED was turned on for 1 hour, and was subsequently turned off. After the LED was turned off, the cells were wrapped in foil to prevent accidental light exposure. Cells were then lysed after the experimental time period.

### HEK Cell Doxycycline Experiment

For the experiment in Figs. 1D-F, cells were stimulated as above and were lysed at the following timepoints: 0 hours (i.e., immediately before adding dox), 0.5 hours after adding dox, 1 hour after adding dox (i.e., immediately before adding ActD), 2 hours after adding dox, 3 hours after adding dox, 4 hours after adding dox, 5 hours after adding dox, 6 hours after adding dox, 7 hours after adding dox, 8 hours after adding dox, 9 hours after adding dox, 10 hours after adding dox, 11 hours after adding dox, and 12 hours after adding dox. Each timepoint consisted of three replicates. On a separate occasion, we collected three replicates at 2.5 hours after adding dox and 4.5 hours after adding dox, and these timepoints functioned as our test timepoints in Supp. Fig. 3. The data from Fig. 1D-F was further used in Fig. 2 and Fig. 3A,B.

### Single Cell Experiments

For all experiments involving single cells, HEK cell cultures were prepared, transfected with 100ng of pAAV-CAG-GFP (Addgene 37825), 200ng of plasmid 147, 100ng of plasmid 116v5, and 100ng of pCMV Tet3G, stimulated with doxycycline, and then silenced with actinomycin D as described above. The three induction conditions were as follows: cells in condition 1 were left in doxycycline for 3 hours prior to lysis; cells in condition 2 were silenced with actinomycin D after 1 hour, and then left for 3 hours, and cells in condition 3 were silenced with actinomycin D after 1 hour and then left for 7 hours. Subsequently, cells were treated with trypsin (Life Technologies, 25300054). Following trypsinization, cells were centrifuged at 850g, washed in cold PBS, and then resuspended in cold PBS. 96 well plates were prepared, with each well containing a solution of 0.2% Triton-X with 2U/uL RNAse inhibitor. Individual cells were sorted into the wells of this wellplate using a Moflo Astrios EQ flow cytometer. Following sorting, the wellplate was sealed, centrifuged, and then placed at −80C overnight.

For the analysis in Fig. 4A-C, cells in condition 2 received plasmid 147B1, while cells in condition 3 received plasmid 147B3. The two populations of cells were mixed following trypsinization and sorted together. By contrast, cells in condition 1 received plasmid 147B1, and were sorted separately from the others.

Library preparation for the single cells proceeded as follows. Plates containing single cells were thawed, and 7uL of nuclease free water was added to the single cells to bring the total volume up to 11uL. Subsequently, reverse transcription was performed using Superscript IV and the SGR-174 RT primers, as in the case of the bulk samples, with the following modifications. RT primers were distributed so that each cell at a given timepoint received an RT primer with a different barcode. In addition, for each timepoint, we performed two no-template RT reactions. Finally, after the 50C step in the Superscript IV protocol, we cooled the samples to 37C and added 20U of Exonuclease 1 (NEB, M0293S) to the reaction to remove excess primers. Samples then remained at 37C for 10 minutes, before proceeding to the 80C heat inactivation step. Following reverse transcription, the RT reactions for all cells and the two no-template controls at a given timepoint were pooled, cleaned with Ampure XP beads at a 1:1 dilution, and were then PCRed using the same protocol as for the bulk samples. Cells were pooled prior to PCR as a way of reducing the number of cycles necessary to achieve amplification. We excluded cells if they received fewer than 150 reads, or if the most common RNA barcode represented fewer than 80% of the total deduplicated reads, which would indicate index swapping between cells.

For experiments involving 10x Single Cell, as in Figs. D-F, cells in the 1 hour condition were induced with doxycycline 1 hour before 10x preparation; cells in the 2 hour condition were induced with doxycycline 2 hours before 10x preparation and silenced with actinomycin D 1 hour later, and left for 1 hour, and cells in the 4 hour condition were induced with doxycycline 4 hours before 10x preparation, silenced with actinomycin D after 1 hour, then left for 3 hours. Cells were then treated with trypsin (Life Technologies, 25300054), spun down at 500g, washed in cold PBS, and then resuspended in cold PBS with 0.04% weight/volume BSA. Cell concentration was counted and the appropriate volume of cells was added to a 10x reaction for a targeted cell recovery of 2000 cells. The reaction was run through a 10x chip according to the 10x Genomics Chromium Single Cell protocol. Following the droplet generation step, the reaction was transferred to a PCR strip-tube and reverse transcription, post-reverse transcription cleanup and cDNA amplification was completed following the 10x Genomics Chromium Single Cell protocol. Following cDNA cleanup, 1-5 uL of the cDNA product was PCRed with Phusion Hot Start Flex DNA Polymerase (New England BioLabs, M0535S), using a barcoded version of SGR-176 and a version of the 10x Genomics cDNA primer that included a P5 addition, with the settings: 30s of 98C denaturation; then 15-20 cycles of 10s denaturation at 98C, 20s annealing at 63C, and then 25s extension at 72C. The rest of the cDNA product was saved for the remainder of the 10x Genomics Chromium Single Cell protocol. The PCR product was then purified with Ampure beads at a 1:1 dilution, then adjusted to a concentration of 4 nM for sequencing. The libraries were sequenced on a MiSeq with the following read structure 28 Read 1, 8 Index 1, 8 Index 2, 200 Read 2.

### Multiplexing

Experiments for Supp. Fig. 4 were conducted as follows. Three wells of 3T3 cells were transfected as described above with 100ng each of pCMV Tet3G, plasmid 133, plasmid 147B1, plasmid 149B3, and plasmid 116v5. Three wells were transfected with 100ng of pCMV Tet3G, 100ng of plasmid 116v5, and 100ng of plasmid 147B1, and 200ng of pAAV-CAG-GFP. Finally, three wells were transfected with 100ng of plasmid 133, 100ng of plasmid 149B3, 100ng of plasmid 116v5, and 200ng of pAAV-CAG-GFP. Subsequently, all 9 wells were irradiated with blue light as described above for 1 hour, and were the placed in darkness. 7 hours after placing the cells in darkness, cells were stimulated with doxycycline as described above. After one hour in doxycycline, cells were lysed.

### Alignment and Edit Counting

The alignment and analysis pipeline for sequencing data is summarized in Supp. Fig. 6. Analysis of sequencing data was performed using custom Matlab code. Briefly, in the case of single cell data, we first performed deduplication using a 9bp UMI on the RT primer (oligo SGR-174). Other datasets were not deduplicated. Reads were then filtered to ensure that they had the minimum necessary read length (67 bases on Read 1, and 184 bases on Read 2). Note that Read 1 was on the RT primer, so Read 1 reads the reverse complement of the RNA sequence. Thus, the expected mutation was A to G on Read 2, and T to C on Read 1. Alignment was performed using all bases that were not As on Read 2, or that were not Ts on Read 1. Reads were considered to be aligned to the template if 95% of the non-A (for Read 2) or non-T (for Read 1) bases matched the template. Furthermore, we required 90% of the bases that were expected to be As on Read 2 or Ts on Read 1 have Q scores greater than 27 (Supp. Fig. 6); reads that failed to achieve this threshold were discarded.

Finally, we required that all reads have at least one edit in Read 1 and at least one edit in Read 2 for analysis. We implemented this requirement because it appeared to eliminate a number of artifacts that we occasionally observed in our data: for example, each well would sometimes have different (large) numbers of RNAs with zero edits or one edit, which would confound attempts to infer timing by gradient descent. As a consequence of this requirement, all of the histograms of edits per RNA presented in this paper appear to show only very few RNAs with fewer than ~12 edits. There are ~12 bases in template A, all of which are on Read 2, that are edited much more quickly than any bases on Read 1. These are of the form UAG, and all form bulges in the RNA secondary structure, which is thought to encourage editing by ADAR. Exclusion of RNAs with zero edits on Read 1 or Read 2 predominantly limits the analysis to RNAs that are already fully edited at all 12 of those As, thus causing all RNAs to have at least 12 edits. RNAs with fewer than 12 edits may be useful for inferring transcriptional dynamics on the order of minutes, which we observed in preliminary experiments not presented here.

### Exponential Model

The exponential model in Supp. Fig. 3 was implemented using custom code in Python, as follows. For each editable position *i* on the template, we assume the likelihood of base *i* being edited follows an exponential distribution with parameter *λ_i_*, to be estimated from the data. Assuming an instantaneous pulse of transcriptional activity at time t=0, the fraction of edited bases for position *i*, *y*_*i*_, can be modelled as the CDF of the exponential distribution:

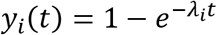

To more accurately capture the experimental setup, we model *y*_*i*_ as an underlying process which is exponential, but with start time uniformly distributed in [0, *t*_*stop*_], where *t*=0 represents when doxycycline is added to the cells and *t*_*stop*_ is the time at which actinomycin D was added to the cells. Specifically, we fit a function of the form

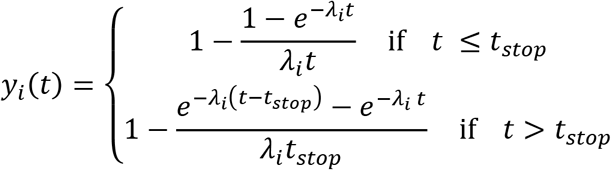

where *t*_*stop*_ was 1hr and *λ*_*i*_ was fit to the data using non-linear least squares. This function was fit for times *t* ≥ 1.5hr, since the editing distributions for earlier timepoints are strongly affected by populations of RNA present prior to doxycycline addition. For the analysis in Supp. Fig. 3, analysis was then performed using only those adenosines for which the R^2^ of the resulting fit was greater than 0.9. We model the total number of edits to the RNA with a Poisson binomial distribution with N trials where N is the total number of editable positions and success probabilities given by *y*_*i*_(*t*) for each position *i*. The probability of having *n* edits at time *t* is given by

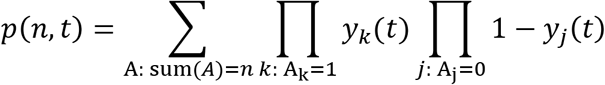

Here, *A* is a binary vector with each entry corresponding to a specific adenosine in the timestamp editing region. *A*_*k*_=1 if adenosine *k* has been edited to inosine, and sum(*A*) counts the total number of edits in A. Time estimates using the exponential model were then made by minimizing the Kullback-Leibler divergence between *p*(*n*,*t*) and the empirical distribution *q*(*n*) over *t*. *p*(*n*,*t*) was calculated in practice via a dynamic programming approach.

For Supp. Fig. 3, the exponential model was calculated using the data from a single replicate of the HEK doxycycline experiment. The distributions in Supp. Fig. 3C show the number of edits per RNA calculated across all bases with R^2^ greater than 0.9 for that replicate, and the Poisson binomial model in Supp. Fig. 3C likewise included the same bases. By contrast, for Supp. Fig. 3D, bases were only retained if they had R^2^ greater than 0.9 in all three replicates from the HEK doxycycline experiment. For this reason, the apparent numbers of edits per RNA are lower in Supp. Fig. 3D than in Supp. Fig. 3C.

### Gradient Descent

The gradient descent in Fig. 2, 3, and Supp. Fig. 5 was implemented using custom code in Matlab. Briefly, the gradient descent algorithm was given an RNA editing distribution (a normalized histogram of edits per RNA), which could either be an experimentally measured distribution (Fig. 2B-D, and Fig. 3C-G) or a simulated distribution (Fig. 3A,B and Supp. Fig. 5). Simulated distributions were generated using data from the experiment in Fig. 1E,F, either by randomly sampling RNAs from a specified timepoint or timepoints from that experiment (as in Fig. 3A,B), or by taking convex combinations of the editing distributions from that experiment (as in Supp. Fig. 5). The gradient descent algorithm was also given a set of “basis vector” histograms, which were obtained by combining the data at each timepoint from all three replicates from the HEK doxycycline experiment. The gradient descent was then initialized by drawing a set of weights from a Dirichlet distribution with all parameters set to unity. The gradient descent minimized the mean squared error (L2 norm) between the input distribution and the convex combination of the basis vectors given by the weights. For each simulated distribution, we performed the gradient descent 1000 times and took the solution that minimized the L2 norm.

To avoid overfitting, whenever an input to the gradient descent was simulated, the distributions used as ‘basis’ functions in the gradient descent were averages of all three biological replicates from the experiment in Fig. 1E,F, while the simulated distributions were always generated from the data associated with a single replicate.

An analysis of the performance of the gradient descent algorithm on random simulated transcriptional programs is presented in Supp. Fig. 5. In general, the gradient descent algorithm succeeded in reproducing the editing histogram with high accuracy (Supp. Fig. 5A). Additionally, the weight vector found by the gradient descent algorithm was on average much closer to the true weight vector than randomly sampled vectors (Supp. Fig. 5B), although this was not always true (Supp. Fig. 5C): because the simulated and approximated editing histograms were generated with different basis distributions, noise present in those basis distributions meant that the true weight vector was not in general the optimal solution for the gradient descent (Supp. Fig. 5D,E).

### Analysis of Two Peaks Experiments

In Fig. 3B,G, a weight distribution generated by the gradient descent algorithm was defined to have two peaks if the sum of the weights on the 1 and 2 hour timepoints and the sum of the weights on the 4 and 5 hour timepoints were both greater than the weight on the 3 hour timepoint. When making Fig. 3B,G, we generated several possible definitions for what would constitute “two peaks,” and obtained qualitatively similar behavior (monotonically increasing identification of two peaks) for all of them. This definition was chosen for presentation arbitrarily.

### Accuracy Metrics

In Fig. 2C, temporal estimation error is calculated by multiplying the distance of each timepoint away from the expected timepoint by the weight assigned to that timepoint, and summing. Thus, for the 3-hour single-induction pulse, if the decoder assigned weights of 0.5 to the 3-hour timepoint and 0.5 to the 5-hour timepoint, the resulting estimation error would be 0.5*0 + 0.5*2=1 hour. This metric is not directly comparable to the 95% confidence interval presented in reference to Fig. 1F, but if the procedure used to generate Fig. 2C is repeated for “sets” consisting of a single RNA, the error obtained is 1.9 ± 0.47 hours (mean ± std, N=12 timepoints), which is significantly less than the 1.49 ± 0.42 hours obtained for sets of 4 RNAs (means differ by >3 standard errors), supporting the statement that the error for 4 RNAs is less than that for a single RNA.

For the arbitrary transcriptional program experiments in Supp. Fig. 5, we calculated the accuracy as the sum of the absolute values of the differences between the assigned and expected weights, divided by 2 to avoid double-counting. Thus, if we expected one timepoint to get 100% of the total weight, and that timepoint instead got 80% of the total weight, then the resulting accuracy would be 80%.

In Fig. 4B, the accuracy is calculated as the mean absolute difference between the single cell estimates and the estimate for the bulk distribution. We calculate the accuracy in this way for the single cells because the ground truth transcriptional program is not known. The single cells stay on ice for up to an hour during processing, and we have not measured the editing kinetics during that time.

