## Supplemental Figures for "Recording the age of RNA with deamination"

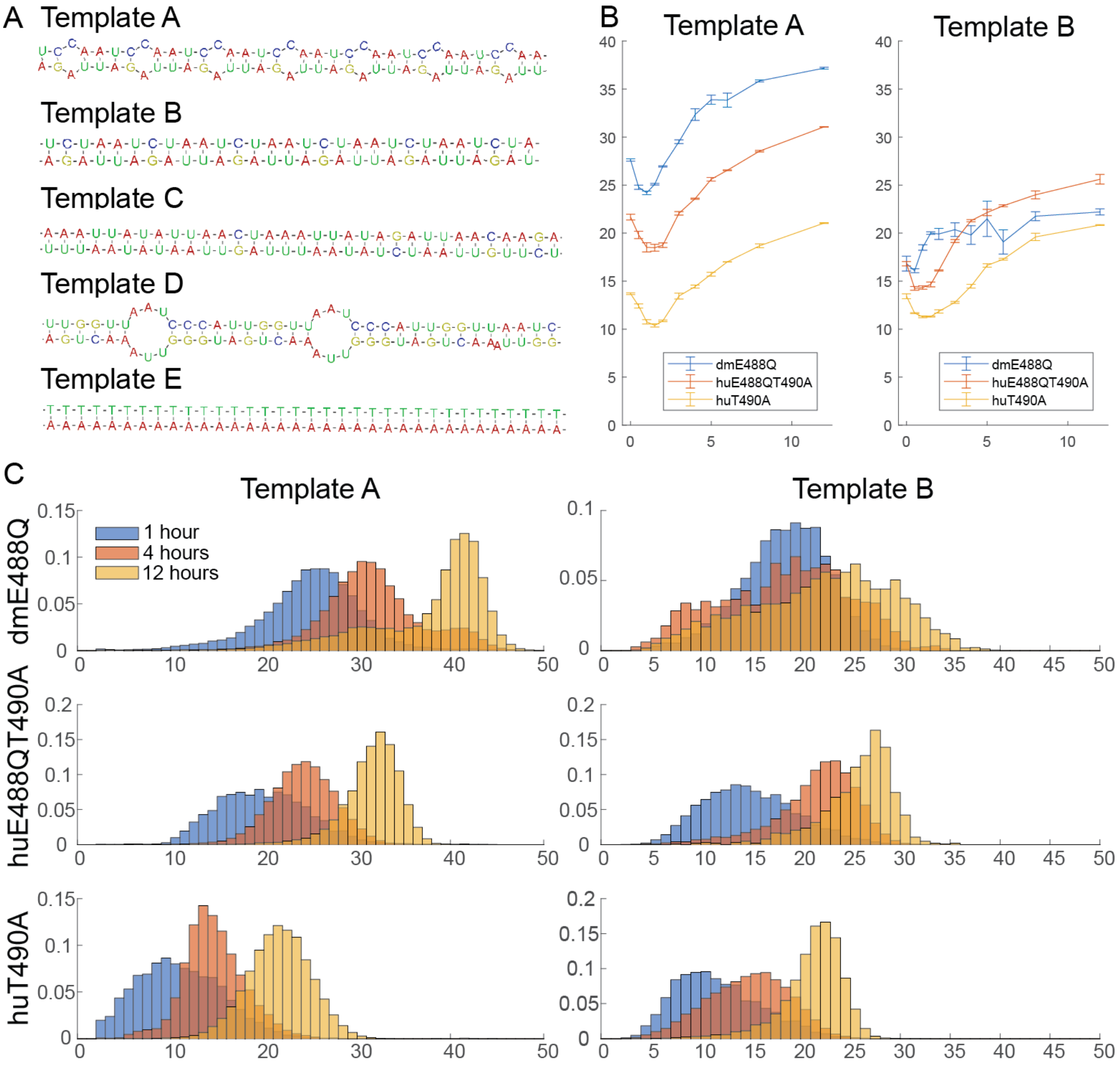


**Supp. Figure 1:** (**A**) Five editing templates that were tested are shown. Of these, only templates A and B showed robust temporal editing that seemed appropriate for the construction of the timestamp system. Notes on the templates are in Supplementary Table 3. (**B**) The mean number of edits per RNA for several different timepoints is shown for three different ADAR variants, and for templates A and B. The protocol used here is identical to that in Fig. 1E. Some combinations, such as dmE488Q with template A, may show greater temporal resolution at short timescales. (**C**) Example editing histograms are shown for three different timepoints, for each combination of the three enzymes and two templates in (B).

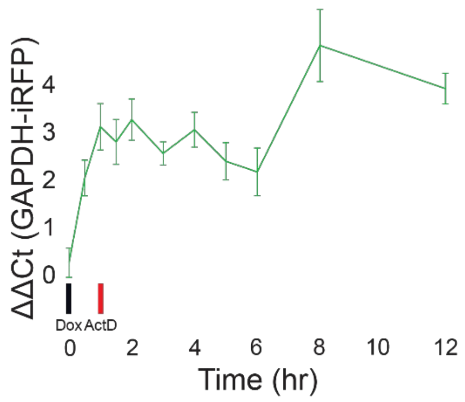

**Supp. Figure 2:** The qPCR for the iRFP transcript, normalized to GAPDH, is shown as a function of time during the experiment in Fig. 1E. Values are normalized to the pre-doxycycline timepoint. Error bars show standard deviation (N=3).

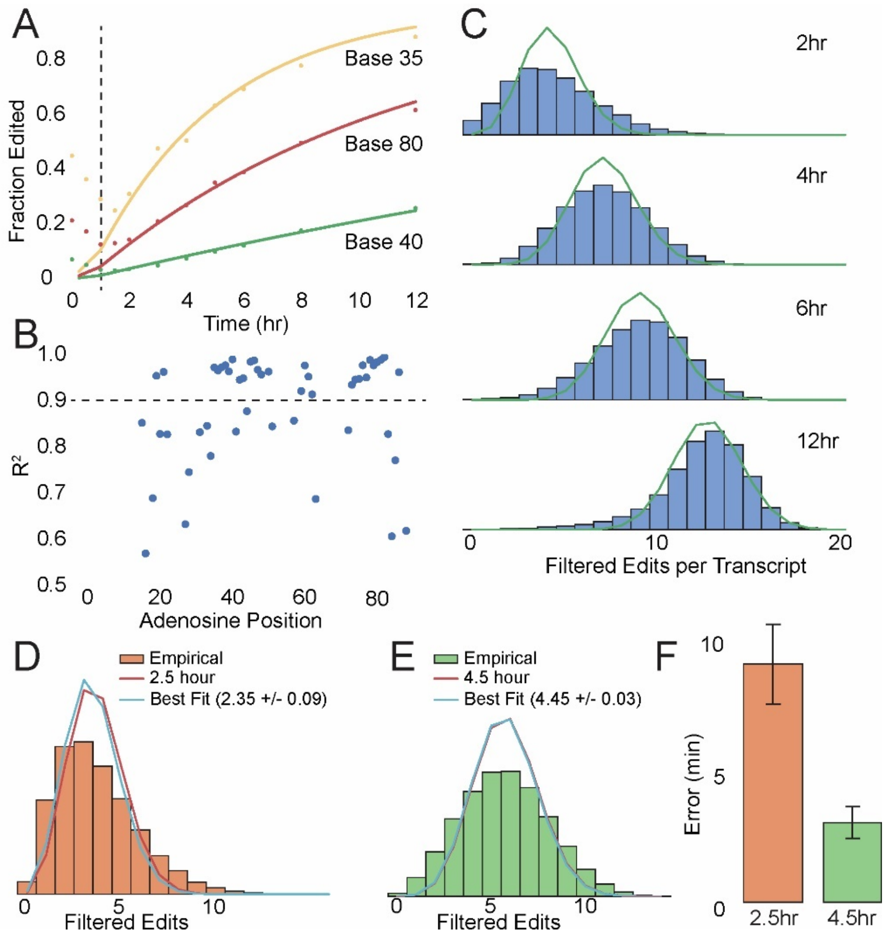


**Supp. Figure 3.** We designed a statistical model to predict the RNA age distribution as a function of time since doxycycline induction. If the adenosines on the timestamp template are edited independently and uniformly in time, then for each adenosine on the timestamp, the fraction of RNAs with adenosines at that site should decrease exponentially with the time since transcription, with a site-specific rate constant that depends on the local sequence context. (**A**) For each adenosine on the timestamp, we fitted an exponential cumulative distribution function (CDF) to the editing fraction over time at that base. The fraction of A to I edits as a function of time is shown for three different bases on the timestamp, data from one replicate of 1E. Best exponential fits are shown. The black dotted line indicates the addition of actinomycin D. (**B**) We found 24 bases which fit well to the model (i.e., for which the value of R2 was greater than 0.9 across all replicates). For the same replicate as in (A), the $R^2$ value of the exponential fit is shown for each base on the transcript. The black dotted line indicates the $R^2 > 0.9$ cutoff used for the exponential model. (**C**) Analyzing only those bases, the distribution of edits per RNAs was well-approximated by a Poisson binomial distribution with a single parameter, t, which represents time since doxycycline was added to the medium (see Methods), with the weights in the Poisson binomial distribution given by the exponential CDFs. The masked editing histograms for four timepoints from the same replicate are shown (only the bases with $R^2 > 0.9$ are included). In green, the Poisson binomial distribution for each timepoint including all the bases with $R^2 > 0.9$ (see Methods). (**D**) We used this Poisson binomial distribution to infer the times of cells induced at 2.5 and 4.5 hours prior to lysis, timepoints that had not been included in the dataset used to fit the exponential CDFs. By minimizing the Kullback-Leibler divergence (which is equivalent to maximizing the likelihood) between the test distributions and the Poisson binomial distribution over t, we inferred that timing of those events to be 2.35hr ± 0.09hr and 4.45hr ± 0.03hr (mean ± s.d., N=3 technical replicates), respectively. In orange, the masked ($R^2$ > 0.9 in all 3 replicates from 1E, see Methods) editing histogram for a single 2.5 hour replicate along with Poisson binomial distribution for 2.5 hours (red line), and the Poisson binomial distribution with least KL divergence from the empirical distribution (blue line). The time estimate is mean ± s.d. (N=3 technical replicates). (**E**) As in (D), but for the 4.5 hour timepoint. (**F**) The mean absolute error is shown for the (D) and (E). Error bars show standard deviation.

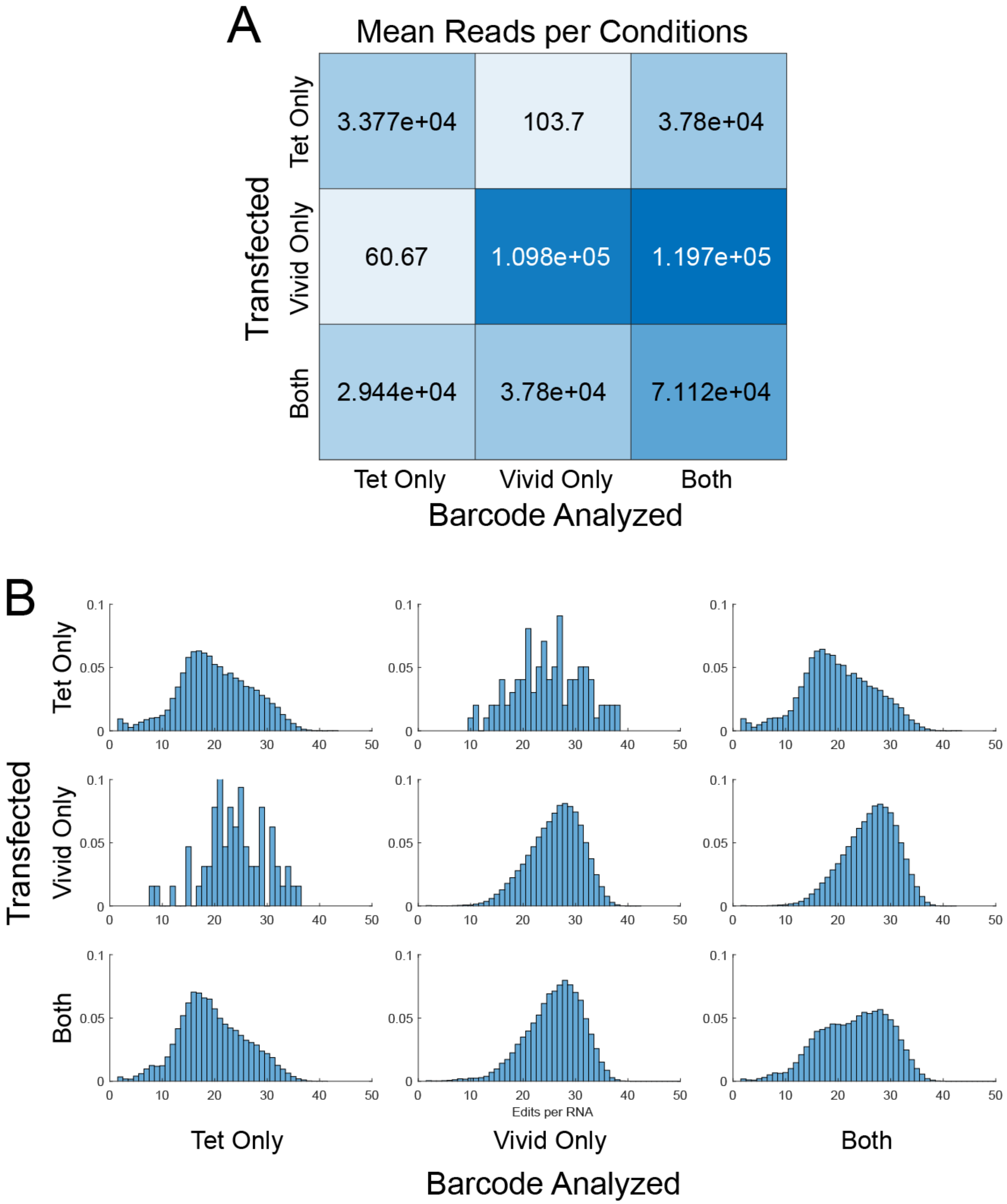


**Supp. Figure 4.** The fact that timestamps work with multiple promoters raises the possibility of recording the activity of multiple promoters simultaneously in a single cell population, and we validated that this is possible using barcoded timestamps responsive to the Tet and Vivid promoters. All editing histograms are normalized to sum to 1. (**A**) For cells transfected with a barcoded TRE-responsive timestamp construct, a barcoded Vivid-responsive timestamp construct, or both, the number of reads for the TRE-responsive timestamp, Vivid-responsive timestamp, or both are shown. When only one timestamp is transfected, only one barcode is detected in significant numbers, confirming that there is minimal crossover between timestamp barcodes. Note that the third column is not the sum of the first and second columns, because it includes barcodes that did not perfectly align to either the Tet or Vivid timestamp barcodes. (**B**) To further confirm the possibility of multiplexing using barcoded timestamps, we analyzed the editing histograms for cells that were transfected with a barcoded TRE-responsive timestamp construct, a barcoded Vivid-responsive timestamp construct, or both. The editing histograms for the Vivid-responsive and TRE-responsive timestamps do not seem to change when the other timestamp is also present, again suggesting that there is minimal cross-talk between barcoded timestamp constructs. All editing histograms are normalized to sum to 1.

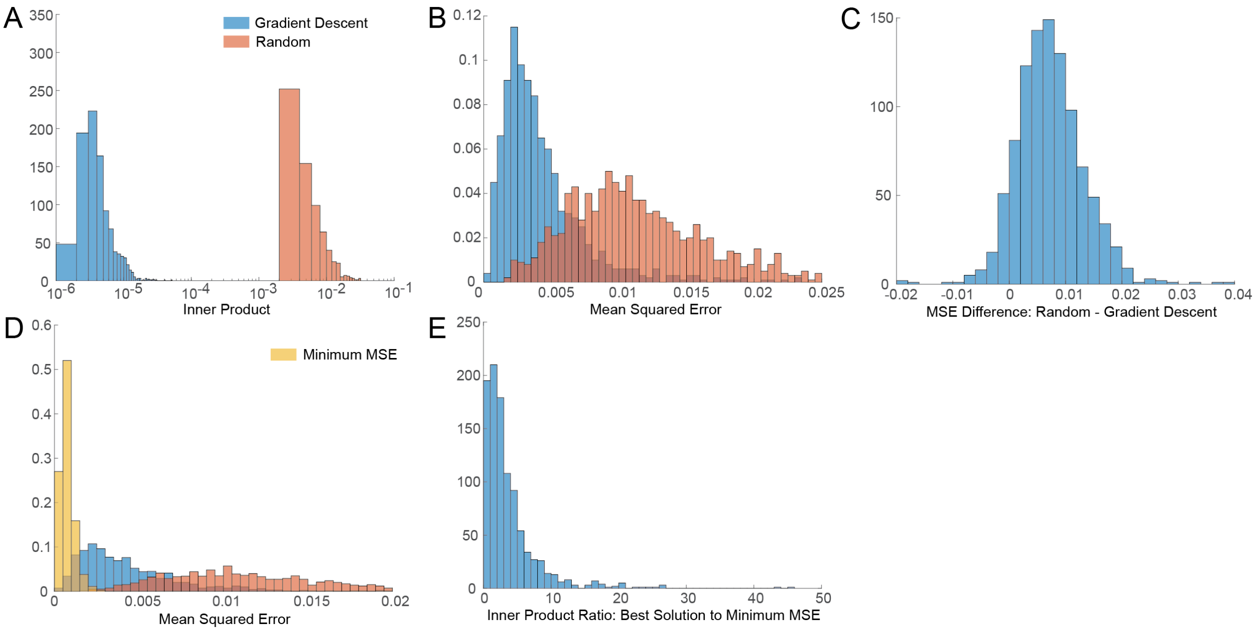


**Supp. Figure 5.** For 1000 randomly generated weight vectors ("simulated vectors"), chosen according to a Dirichlet distribution with uniform weights, we used gradient descent to find the approximation ("approximated vectors") that minimized the L2 norm ("inner product") between the RNA editing distribution corresponding to the simulated vectors ("simulated distributions") and the RNA editing distribution corresponding to the approximated vectors ("approximated distributions"). We refer to the L2 norm between the distributions as the inner product to distinguish it from the L2 norm between the vectors, which we refer to as the mean squared error (MSE). (**A**) The inner product between simulated distributions and approximated distributions is shown in blue. By contrast, the inner product between simulated distributions and other random distributions is shown in orange. (**B**) The mean squared error between the simulated vectors and approximated vectors is shown in blue. By contrast, the inner product between the simulated distributions and other random distributions is shown in orange. Note that a substantial number of random weight vectors have lower mean squared error than the approximated vectors. This is possible because the noise in the basis distribution set used to generate the approximated distributions from the approximated vectors is different from the noise in the basis distribution set used to generate the simulated distributions from the simulated vectors, so the minimum of inner product between the simulated and approximated distributions is not always the same as the minimum of the MSE between the simulated and approximated vectors. (**C**) Another visualization of (B). For each simulated vector, we calculated both an approximated vector and a random vector. The difference in MSE between the approximated and random vectors is shown. Negative values correspond to test vectors for which the associated random vector was a better approximation to the simulated vector than the approximated vector. (**D**) Blue and orange bars are the same as in (B). Yellow bars correspond to the minimum MSE among all of the solutions found by gradient descent for a given test vector, indicating that the inner product minima found by the gradient descent are not in general minima of the MSE. (**E**) The difference in the inner product between the solutions with the minimum MSE found by gradient descent, and the solutions with the minimum inner product, as a fraction of the minimum inner product. The solutions with the minimum MSE discovered by gradient descent often have inner products several fold higher than the solution with the minimum inner product.

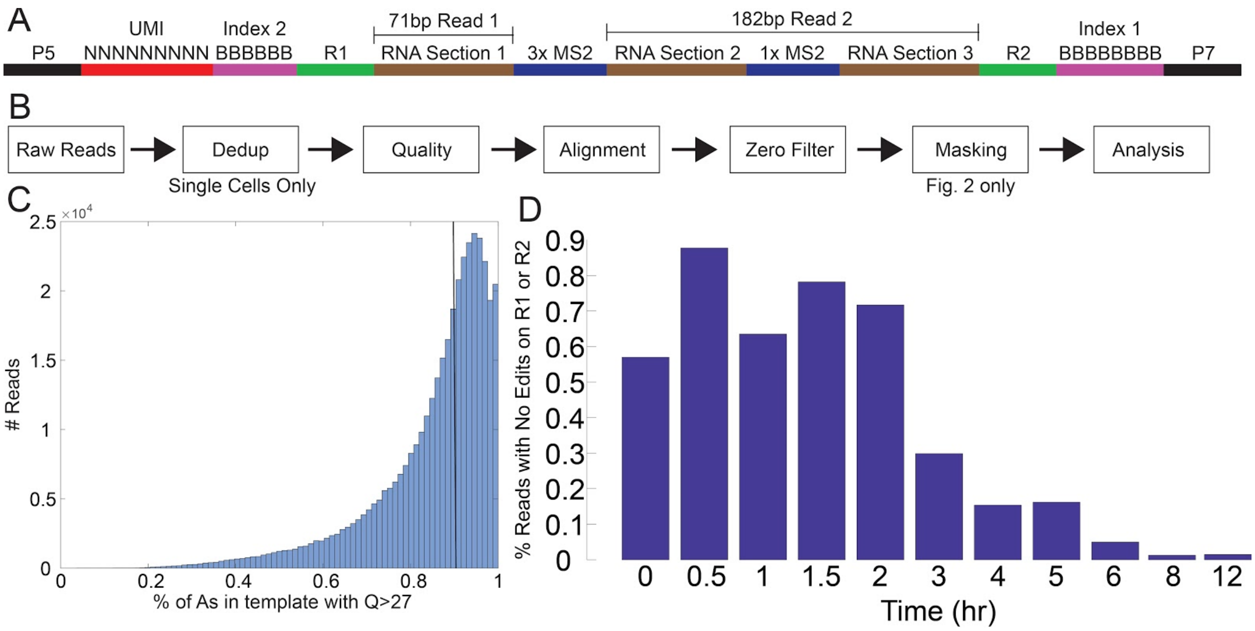


**Supp. Fig. 6:** (**A**) The read structure of the timestamp is shown. (**B**) A schematic of the analysis pipeline is shown. See Methods. (**C**) For one replicate from the experiment in Fig. 1E, a histogram of the number of reads with a given percentage of As with Q score >27 is shown. This includes all sites that are As on the timestamp template, i.e., it also counts Gs that are read at positions that are A on the template. The black line indicates the 90% cutoff, which was applied to all analysis. (**D**) For one replicate from the experiment in Fig. 1E, the percentage of reads having no edits in either R1 or R2 is shown as a function of time. These reads were excluded from analysis, except where otherwise stated in Fig. S2.

**Supplementary Table 1:** List of plasmids used in this study. This list excludes pCMV Tet3G, which is available commercially from Clontech.

| Num | Name | Description | Used in |
|---|---|---|---|
| 116v1 | pAAV-Ef1a-MCP-dmADARE488Q | Fusion of MS2 coat protein to Drosophila ADAR E488Q, under Ef1a promoter, with WPRE | Supp. Fig 1B,C |
| 116v5 | pAAV-Ef1a-MCP-huADARE488QT490A | As with 116v1, but Human ADAR2 E488QT490A | All figures |
| 116v6 | pAAV-Ef1a-MCP-huADART490A | As with 116v1, but Human ADAR2 T490A | Supp. Fig. 1B,C |
| 133 | pcDNA3.1-GAVPO | GAVPO (VIVID transactivator) expressed under the CMV promoter in the pcDNA3.1 backbone. | Fig. 3C-G, Supp. Fig. 4 |
| 147B1 | pTRE3G-iRFP-B1-timestamp_A | Timestamp Template A inserted into the 3' UTR of iRFP between a bActin Zipcode element and a WPRE element, in the pTRE3G backbone, with RNA barcode TGC. Also includes a xrRNA element in the 5' UTR. | All Figures |
| 148B1 | pTRE3G-iRFP-B1-timestamp_B | Same as 147B1, but with RNA Template B. | Supp. Fig. 1 |
| 149B3 | pLenti-5xUASG-iRFP-B3-timestamp-A | timestamp Template A inserted into the 3' UTR of iRFP between a bActin Zipcode element and a WPRE element, in a second generation lentiviral backbone with the Vivid promoter, with RNA barcode CTG. Also includes a xrRNA element in the 5' UTR. | Fig. 3C-G, Supp. Fig. 4 |

**Supplementary Table 2:** List of oligos used in this study

| Name | Description | Sequence |
|---|---|---|
| SGR-174B-1 | Barcoded RT Primer with 3bp barcode | AATGATACGGCGACCACCGAGATCTACACNNNNNNNNNNNN CCT GCG AGG CCC GCATCTTTCACAAATTTTGTAATCCAGAGG |
| SGR-174B-2 | “” | AATGATACGGCGACCACCGAGATCTACACNNNNNNNNNNNN GAG GCG AGG CCC GCATCTTTCACAAATTTTGTAATCCAGAGG |
| SGR-174B-3 | “” | AATGATACGGCGACCACCGAGATCTACACNNNNNNNNNNNN TTA GCG AGG CCC GCATCTTTCACAAATTTTGTAATCCAGAGG |
| SGR-174B-4 | “” | AATGATACGGCGACCACCGAGATCTACACNNNNNNNNNNNN AGC GCG AGG CCC GCATCTTTCACAAATTTTGTAATCCAGAGG |
| SGR-174B-5 | “” | AATGATACGGCGACCACCGAGATCTACACNNNNNNNNNNNN AAT GCG AGG CCC GCATCTTTCACAAATTTTGTAATCCAGAGG |
| SGR-174B-6 | “” | AATGATACGGCGACCACCGAGATCTACACNNNNNNNNNNNN CAA GCG AGG CCC GCATCTTTCACAAATTTTGTAATCCAGAGG |
| SGR-174B-7 | Barcoded RT primer with 6 base barcode | AATGATACGGCGACCACCGAGATCTACACNNNNNNNNNAGTGT CGCG AGG CCC GCATCTTTCACAAATTTTGTAATCCAGAGG |
| SGR-174B-8 | “” | AATGATACGGCGACCACCGAGATCTACACNNNNNNNNNTATCC GGCG AGG CCC GCATCTTTCACAAATTTTGTAATCCAGAGG |
| SGR-174B-9 | “” | AATGATACGGCGACCACCGAGATCTACACNNNNNNNNNCATTT GGCG AGG CCC GCATCTTTCACAAATTTTGTAATCCAGAGG |
| SGR-174B-10 | “” | AATGATACGGCGACCACCGAGATCTACACNNNNNNNNNATGCT AGCG AGG CCC GCATCTTTCACAAATTTTGTAATCCAGAGG |
| SGR-174B-11 | “” | AATGATACGGCGACCACCGAGATCTACACNNNNNNNNNCCGTG GGCG AGG CCC GCATCTTTCACAAATTTTGTAATCCAGAGG |
| SGR-174B-12 | “” | AATGATACGGCGACCACCGAGATCTACACNNNNNNNNNATGAG TGCG AGG CCC GCATCTTTCACAAATTTTGTAATCCAGAGG |
| SGR-174B-13 | “” | AATGATACGGCGACCACCGAGATCTACACNNNNNNNNNCGAGC AGCG AGG CCC GCATCTTTCACAAATTTTGTAATCCAGAGG |
| SGR-174B-14 | “” | AATGATACGGCGACCACCGAGATCTACACNNNNNNNNNCGCGG CGCG AGG CCC GCATCTTTCACAAATTTTGTAATCCAGAGG |
| SGR-174B-15 | “” | AATGATACGGCGACCACCGAGATCTACACNNNNNNNNNACTTA TGCG AGG CCC GCATCTTTCACAAATTTTGTAATCCAGAGG |
| SGR-174B-16 | “” | AATGATACGGCGACCACCGAGATCTACACNNNNNNNNNTGCAT GGCG AGG CCC GCATCTTTCACAAATTTTGTAATCCAGAGG |
| SGR-174B-17 | “” | AATGATACGGCGACCACCGAGATCTACACNNNNNNNNNAGTAG GGCG AGG CCC GCATCTTTCACAAATTTTGTAATCCAGAGG |
| SGR-174B-18 | “” | AATGATACGGCGACCACCGAGATCTACACNNNNNNNNNGTTGA CGCG AGG CCC GCATCTTTCACAAATTTTGTAATCCAGAGG |
| SGR-174B-19 | “” | AATGATACGGCGACCACCGAGATCTACACNNNNNNNNNTATCA CGCG AGG CCC GCATCTTTCACAAATTTTGTAATCCAGAGG |

| | | |
|---|---|---|
| SGR-174B-20 | “” | AATGATACGGCGACCACCGAGATCTACACNNNNNNNNNCCCTAGGCG AGG CCC GCATCTTTCACAAATTTTGTAATCCAGAGG |
| SGR-174B-21 | “” | AATGATACGGCGACCACCGAGATCTACACNNNNNNNNNGCCCGTGCG AGG CCC GCATCTTTCACAAATTTTGTAATCCAGAGG |
| SGR-174B-22 | “” | AATGATACGGCGACCACCGAGATCTACACNNNNNNNNNTTCCCGGCG AGG CCC GCATCTTTCACAAATTTTGTAATCCAGAGG |
| SGR-174B-23 | “” | AATGATACGGCGACCACCGAGATCTACACNNNNNNNNNCATATAGCG AGG CCC GCATCTTTCACAAATTTTGTAATCCAGAGG |
| SGR-174B-24 | “” | AATGATACGGCGACCACCGAGATCTACACNNNNNNNNNAACGCCGCG AGG CCC GCATCTTTCACAAATTTTGTAATCCAGAGG |
| SGR-174B-25 | “” | AATGATACGGCGACCACCGAGATCTACACNNNNNNNNNAGGTTGGCG AGG CCC GCATCTTTCACAAATTTTGTAATCCAGAGG |
| SGR-174B-26 | “” | AATGATACGGCGACCACCGAGATCTACACNNNNNNNNNTCAATAGCG AGG CCC GCATCTTTCACAAATTTTGTAATCCAGAGG |
| SGR-175 | Custom Read 1 | GCG AGG CCC GCA TCT TTC ACA AAT TTT GTA ATC CAG AGG |
| SGR-175-RC | Custom Index 2 | CCTCTGGATTACAAAATTTGTGAAAGATGCGGGCCTCGC |
| SGR-176 | Barcoded PCR primer | CAAGCAGAAGACGGCATACGAGAT ACTGGTCA AAG TTA CTA TCG AAATGCCCTGAGTCCACCCCGG |
| SGR-176-2 | “” | CAAGCAGAAGACGGCATACGAGAT GTGTTCGT AAG TTA CTA TCG AAATGCCCTGAGTCCACCCCGG |
| SGR-176-3 | “” | CAAGCAGAAGACGGCATACGAGAT TAACTGTT AAG TTA CTA TCG AAATGCCCTGAGTCCACCCCGG |
| SGR-176-4 | “” | CAAGCAGAAGACGGCATACGAGAT GATTGGTG AAG TTA CTA TCG AAATGCCCTGAGTCCACCCCGG |
| SGR-176-5 | “” | CAAGCAGAAGACGGCATACGAGAT GGAGAGAG AAG TTA CTA TCG AAATGCCCTGAGTCCACCCCGG |
| SGR-176-6 | “” | CAAGCAGAAGACGGCATACGAGAT TGAGCGAT AAG TTA CTA TCG AAATGCCCTGAGTCCACCCCGG |
| SGR-176-7 | “” | CAAGCAGAAGACGGCATACGAGAT CCTCCGTT AAG TTA CTA TCG AAATGCCCTGAGTCCACCCCGG |
| SGR-176-8 | “” | CAAGCAGAAGACGGCATACGAGAT AACATATT AAG TTA CTA TCG AAATGCCCTGAGTCCACCCCGG |
| SGR-176-9 | “” | CAAGCAGAAGACGGCATACGAGAT CTTACGTA AAG TTA CTA TCG AAATGCCCTGAGTCCACCCCGG |
| SGR-176-10 | “” | CAAGCAGAAGACGGCATACGAGAT TGACGTAG AAG TTA CTA TCG AAATGCCCTGAGTCCACCCCGG |
| SGR-176-11 | “” | CAAGCAGAAGACGGCATACGAGAT CTATGTAT AAG TTA CTA TCG AAATGCCCTGAGTCCACCCCGG |
| SGR-176-12 | “” | CAAGCAGAAGACGGCATACGAGAT TTTGCAGA AAG TTA CTA TCG AAATGCCCTGAGTCCACCCCGG |

| | | |
|---|---|---|
| SGR-176-13 | “” | CAAGCAGAAGACGGCATACGAGAT GGTAGCGA AAG TTA CTA TCG AAATGCCCTGAGTCCACCCCGG |
| SGR-176-14 | “” | CAAGCAGAAGACGGCATACGAGAT ACGGGTTT AAG TTA CTA TCG AAATGCCCTGAGTCCACCCCGG |
| SGR-176-15 | “” | CAAGCAGAAGACGGCATACGAGAT TAAACCTC AAG TTA CTA TCG AAATGCCCTGAGTCCACCCCGG |
| SGR-176-16 | “” | CAAGCAGAAGACGGCATACGAGAT GAGAACTG AAG TTA CTA TCG AAATGCCCTGAGTCCACCCCGG |
| SGR-176-15 | “” | CAAGCAGAAGACGGCATACGAGAT GGTTTGAT AAG TTA CTA TCG AAATGCCCTGAGTCCACCCCGG |
| SGR-176-18 | “” | CAAGCAGAAGACGGCATACGAGAT TAGATTAT AAG TTA CTA TCG AAATGCCCTGAGTCCACCCCGG |
| SGR-176-19 | “” | CAAGCAGAAGACGGCATACGAGAT AAGGTTAG AAG TTA CTA TCG AAATGCCCTGAGTCCACCCCGG |
| SGR-176-20 | “” | CAAGCAGAAGACGGCATACGAGAT CCGAAAAT AAG TTA CTA TCG AAATGCCCTGAGTCCACCCCGG |
| SGR-177 | Custom Read 2 | AAG TTA CTA TCG AAA TGC CCT GAG TCC ACC CCG G |
| SGR-177-RC | Custom Index 1 | CCGGGGTGGACTCAGGGCATTTCGATAGTAACTT |

**Supplementary Table 3:** List of RNA editing templates used in this study.

The following sequences are the sequences that were analyzed for RNA editing. Notes are supplied as a courtesy to follow-on studies, and no representations are made as to their accuracy or reproducibility.

| | **Sequence** | **Notes** |
|---|---|---|
| **A_Short** | AGTACGCGTTAGATTAGATTAGATTAGATTA GATTAGATTAGAAAAATTAATACGTACACC ATCAGGGTACGTCTCAGACACCATCAGGGT CTGTCTGGTACAGCATCAGCGTACCATATAT TTTTTCCAATCCAATCCAATCCAATCCAATC CAATCCAAATAGATCCTAATCA | |
| **A** | TTAGATTAGATTAGATTAGATTAGATTAGAT TAGAAAAATTAATATACGTACACCATCAGG GTACGTCATATATTTTTTCCAATCCAATCCA ATCCAATCCAATCCAATCCAATACGCGTTAG ATTAGATTAGATTAGATTAGATTAGATTAGA AAAATTAATACGTACACCATCAGGGTACGT CTCAGACACCATCAGGGTCTGTCTGGTACAG CATCAGCGTACCATATATTTTTTCCAATCCA ATCCAATCCAATCCAATCCAATCCAAATAGA TCCTAATCA | |
| **B_Short** | AGTACGCGTTAGATTAGATTAGATTAGATTA GATTAGATTAGAAAAATTAATACGTACACC ATCAGGGTACGTCTCAGACACCATCAGGGT CTGTCTGGTACAGCATCAGCGTACCATATAT TTTTTCTAATCTAATCTAATCTAATCTAATCT AATCTAAATAGATCCTAATCA | |
| **B** | TTAGATTAGATTAGATTAGATTAGATTAGAT TAGAAAAATTAATATACGTACACCATCAGG GTACGTCATATATTTTTTCTAATCTAATCTAA TCTAATCTAATCTAATCTAAACGCGTTAGAT TAGATTAGATTAGATTAGATTAGATTAGAAA AATTAATACGTACACCATCAGGGTACGTCTC AGACACCATCAGGGTCTGTCTGGTACAGCAT CAGCGTACCATATATTTTTTCTAATCTAATCT AATCTAATCTAATCTAATCTAAATAGATCCT AATCA | |
| **C** | AGTACGCGTTAAATTATATTAACTAAATTAT AGATTAACAAGAATATTAAATACGTACACC ATCAGGGTACGTCTCAGACACCATCAGGGT CTGTCTGGTACAGCATCAGCGTACCTATTTA ATATTCTTGTTAATCTATAATTTAGTTAATAT AATTTAAATAGATCCTAATCA | This template shows significant background editing by endogenous ADAR enzymes, even in the absence of trans-expression of ADAR. It also showed extremely rapid editing on a timescale of single minutes in the presence of blue light, |

| | | |
|---|---|---|
| | | when MCP-Cry2 and CIBN-dmADARE488Q were co-expressed. |
| **D** | AGTACGCGATTGGTTAATCCCATTGGTTAATCCCATTGGTTAATCCCTTAATACGTACACCATCAGGGTACGTCTCAGACACCATCAGGGTCTGTCTGGTACAGCATCAGCGTACCATATATGGGTTAAACTGATGGGTTAAACTGATGGGTTAAACTGATATAGATCCTAATCA | Editing on this template showed significant sensitivity to the identity of the N-terminal fusion. MCP-ADAR was able to edit this template, whereas other ADAR enzymes, like a CIBN-ADAR fusion, were unable. |
| **E** | AGTACGCGAAAAAAAAAAAAAAAAAAAAAAAAAAAAAAAAAAAAAAAAAAAAACGTACACCATCAGGGTACGTCTCAGACACCATCAGGGTCTGTCTGGTACAGCATCAGCGTACCTTTTTTTTTTTTTTTTTTTTTTTTTTTTTTTTTTTTTTTTTTTTTATAGATCCTAATCA | This template was always severely underrepresented in sequencing, either due to difficulties with expression, amplification, or alignment. |
